# High-throughput bactericidal assays for monoclonal antibody screening against antimicrobial resistant *Neisseria gonorrhoeae*

**DOI:** 10.1101/2023.05.10.540186

**Authors:** Samuele Stazzoni, Marco Troisi, Valentina Abbiento, Claudia Sala, Emanuele Andreano, Rino Rappuoli

## Abstract

*Neisseria gonorrhoeae* (gonococcus) is an obligate human pathogen and the etiological agent of the sexually transmitted disease gonorrhea. The rapid rise in gonococcal resistance to all currently available antimicrobials has become a significant public health burden and the need to develop novel therapeutic and prophylactic tools is now a global priority. While high-throughput screening methods allowed rapid discovery of extremely potent monoclonal antibodies (mAbs) against viral pathogens, the field of bacteriology suffers from the lack of assays that allow efficient screening of large panels of samples. To address this point, we developed luminescence-based (L-ABA) and resazurin-based (R-ABA) antibody bactericidal assays that measure *N. gonorrhoeae* metabolic activity as a proxy of bacterial viability. Both L-ABA and R-ABA are applicable on the large scale for the rapid identification of bactericidal antibodies and were validated by conventional methods. Implementation of these approaches will be instrumental to the development of new medications and vaccines against *N. gonorrhoeae* and other bacterial pathogens to support the fight against antimicrobial resistance.

## INTRODUCTION

Gonorrhea is a sexually transmitted disease with an estimated incidence of 86.9 million cases globally per annum (1). It is caused by the host-adapted human pathogen *N. gonorrhoeae* (2), which is listed as a high priority pathogen for research into novel treatments by the World Health Organization (WHO) because of its ability to quickly develop resistance to antibiotics (3,4). Recently emerged gonococcus strains showed resistance to most currently available antibiotics including ceftriaxone which represents the last remaining option for first-line treatment (5,6). Despite efforts over many decades, no vaccine or specific treatment have yet been successfully developed to counter gonococcal infections and novel strategies need to be implemented to achieve this goal (7). Human monoclonal antibodies (mAbs) have shown great potential in the fight against infectious diseases, especially against viral pathogens. Indeed, mAbs are extremely effective and specific toward their targets, and nowadays can be developed faster than any other drug. The coronavirus disease 2019 (COVID-19) pandemic case has highlighted their great potential and the possibility to bring mAbs from bench to bedside in less than 6 months (8). This milestone was achieved thanks to the unprecedented technological and methodological advancement of the last two decades in the field of B cell analyses and mAb screening. Indeed, this scenario has permitted to interrogate the human antibody repertoire and to quickly screen hundreds of thousands of antibodies to select the optimal candidates for clinical development. Conversely, the field of mAb screening and selection against bacterial pathogens is hampered by the lack of efficient screening assays. The serum bactericidal assay (SBA), conventionally used to evaluate the functional activity of sera or mAbs *in vitro*, is a long and lab-intensive method that relies on counting the number of colonies of survived bacteria on agar plates (**Fig. 1C**) (9). While this approach has been extensively used as an *in vitro* surrogate assay to evaluate the immunogenicity of bacterial vaccines against cholera (10) and meningococcal disease (11), it is not suitable for high-throughput mAb screening. To respond to this unmet need, we herein report the development of two high-throughput (HT) assays named luminescence-based (L-ABA) and resazurin-based (R-ABA) antibody bactericidal assays (**Fig. 1A-B**), for rapid screening and identification of antibodies against gonococcus and AMR bacteria.

**Fig. 1:**
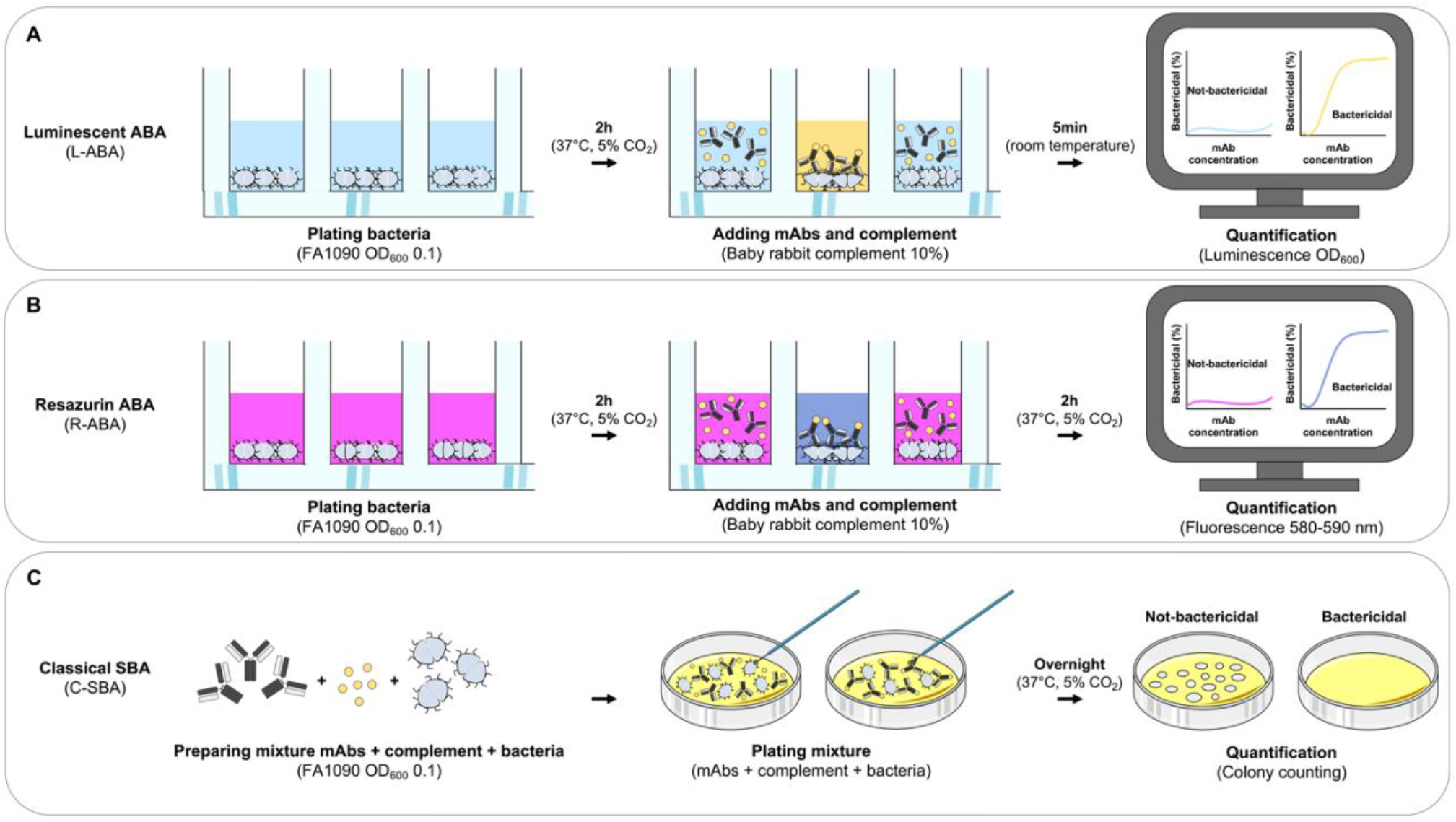
Workflow for monoclonal antibody based high-throughput serum bactericidal assays. (A-C) schematic representation of the workflows implemented for the L-ABA (A), R-ABA (B) and C-SBA C).

## MATERIALS AND METHODS

### Bacterial strain and preparation

*N. gonorrhoeae* strain FA1090 (ATCC 700825), was used in this study. Fresh cultures of bacteria were prepared from frozen stocks by streaking onto gonococcal agar (GCA) consisting of gonococcal (GC) agar base supplemented with 1% v/v IsoVitaleX (BD Biosciences, Franklin Lakes, NJ, USA). On the following day, bacteria were grown in GC liquid medium at 37°C, 5% CO_2_ starting from an optical density at 600 nanometers (OD_600_)0.1 until mid-log phase cultures, i.e., OD_600_ 0.5.

### Source of complement

Baby rabbit complement (BRC) (Cedarlane) was used as a source of complement for serum sensitivity and for all ABA assays. For optimized ABA assays, BRC was diluted to 10% v/v for use as a complement source, which is within the recommended range of 10%–20% v/v for gonococcal bactericidal assays (12). For negative controls with heat-inactivated complement (hiBRC), BRC was heated at 56°C for 1h.

### Complement toxicity assay

*N. gonorrhoeae* was grown to mid-log phase (OD_600_ = 0.5) as described above. Bacterial cultures were then directly diluted in sterile Dulbecco’s phosphate-buffered saline (DPBS) supplemented with 2% fetal bovine serum (FBS) (Hyclone) and 0.1% glucose (from now on named as ABA buffer). To assess the optimal OD_600_, different dilutions were used. Specifically, starting from cultures at OD_600_ 0.5, bacteria were diluted 1:5, 1:10 and 1:100 to obtain final OD_600_ 0.1, 0.01 and 0.001, respectively. Reactions were performed by independently seeding 10 µL of diluted bacteria (ABA buffer) in the presence of 5, 10, 20 or 40 % v/v of BRC (or hiBRC in negative controls) in a final volume of 50 µL. Reactions were incubated for 1 or 2 h at 37°C, 5% CO_2_ in sterile round bottom 96-well plates. After the incubation, plates were centrifuged for 8 min at 4,500 x g. The supernatant was discarded to remove dead or lysed bacteria and the pellets were re-suspended in 30 µL of DPBS. This suspension was transferred into an opaque white 96-well flat bottom plate (Optiplate 96, Perkin Elmer) and 30 µL of BacTiter-Glo reagent (Promega, US) were added to each well. The reaction was incubated for 5 min. After incubation, the luminescence signal was read using a Varioskan™ LUX multimode microplate reader (Thermo Fisher Scientific, Waltham, MA, USA).

### L-ABA assay with purified mAbs

Mid-log phase bacteria (OD_600_ 0.5) were diluted 1:10 in ABA buffer to prepare a 5x working stock. Reactions were performed by seeding 10 µL of 5x bacterial stock with serial dilutions of the purified mAb 2C7 (or a negative control) in ABA buffer and in the presence of 10% v/v BRC (or hiBRC in negative controls) in a final volume of 50 µL for each well. Reactions were incubated for 2 h at 37°C, 5% CO_2_ in sterile round bottom 96-well plates. After the incubation, plates were centrifuged for 8 min at 4,500 x g. The supernatant was discarded to remove dead or lysed bacteria and the pellets were re-suspended in 30 µL of DPBS. This suspension was transferred into an opaque white 96-well flat bottom plate (Optiplate 96, Perkin Elmer) and 30 µL of BacTiter-Glo reagent (Promega, US) were added to each well. The reaction was incubated for 5 min before the luminescence signal was read using a Varioskan™ LUX multimode microplate reader (Thermo Fisher Scientific, Waltham, MA, USA). Acquired data were analyzed using GraphPad Prism version 9.4.1 by fitting the data to a sigmoidal dose-response curve to calculate the IC_50_.

### R-ABA assay with purified mAbs

*N. gonorrhoeae* was grown to mid-log phase (OD_600_ 0.5) as described above. Assay plates were then prepared as described above for the L-ABA, with the only difference that black 96-well plates (Culturplate 96, Perkin Elmer) were used, instead of the round bottom ones. Reactions were incubated for 2 h at 37°C, 5% CO_2_. Then, 10 µL of 0.025% resazurin solution in sterile distilled water were added to each well. The reactions were then further incubated for 2 h at 37°C, 5% CO_2_. At the end of the incubation, fluorescence signals were measured by a Varioskan™ LUX multimode microplate reader (Thermo Fisher, Waltham, MA, USA) using 560 nm for excitation and 590 nm for emission. After the measurement, the assay plate was kept at 37°C overnight to allow complete conversion of resazurin to resorufin in the wells containing live bacteria. On the following day, the plate was observed by eye and pictures were taken.

### Transcriptionally active PCR (TAP-PCR) for 2C7 transfection

To obtain PCR fragments that could be directly used to transfect cells and express mAb 2C7, TAP-PCR reactions of the heavy/light chain fragments were performed in replicates, as previously described (13). Briefly, the appropriate vectors of heavy and light chains, carrying the variable region of the 2C7 mAb, were used as templates. Reaction was performed using 0.25 μL of Q5 polymerase (NEB), 5 μL of GC Enhancer (NEB), 5 μL of 5X buffer,10 mM dNTPs, 0.125 µL of 100μM forward/reverse primers, and 3 µL of template (i.e. vectors carrying heavy and light chains) using the following PCR program: 98°C/2′, 35 cycles composed of 98°C/10″, 61°C/20″, 72°C/1′, and 72°C/5′. TAP-PCR products were purified and quantified by Qubit Fluorometric Quantitation assay (Invitrogen) and used for transient transfection in the Expi293F cell line following manufacturing instructions (Thermo Fisher Scientific).

### 96 well/plate small-scale transfection

Expi293F cells (Thermo Fisher Scientific) were handled as recommended by the manufacturer. Briefly, on the day before the transfection, cells were counted and adjusted at a concentration of 2.5 × 10^6^ cells/mL in Expi293 transfection medium. The day after, cells were counted, centrifuged at 310 x g for 8 minutes and then resuspended to achieve a concentration of 2.5 ×10^6^ cells/mL. 0.8 mL/well of this cell suspension were then added into Deep Well 96-well plates (Eppendorf) and kept at 37°C, 8% CO_2_, in a plate shaker set at 1,000 rpm until cell transfection. Small scale transfection was then carried out according to the manufacturer protocol with the layout described below. Cells in rows A-D were transfected with 300 ng of heavy chain TAP product + 600 ng of light chain TAP product to obtain 2C7-containing supernatants (positive controls). Cells in rows E-H were transfected with lipofectamine-Optimem only, as negative controls (mock transfection of Expi293F cells). After transfection, cells were incubated shaking at 1,000 rpm for 6 days and then centrifuged to collect the supernatants that were used in the HT-ABA assays.

### L-ABA for high-throughput screening of mAb supernatants

Up to 96 different supernatants can be assayed within each plate, using the same setup described above. To evaluate the applicability of this assay for bactericidal screenings, a round-bottom 96-well plate was prepared as follows:

- Rows A-D: each well contained 10 µL supernatant from Expi293F cells transfected with 2C7 (positive control, 1:5 dilution), 5 µL BRC (10% v/v), 10 µL of bacteria (OD_600_ 0.05), 25 µL of ABA buffer to reach a final volume of 50 µL.
- Rows E-H: each well contained 10 µL supernatant of mock Expi293F cells (negative control, 1:5 dilution), 5 µL BRC (10% v/v), 10 µL of bacteria (OD_600_ 0.05), 25 µL of ABA buffer to reach a final volume of 50 µL.

After 2h incubation, the plates were centrifuged and then processed as described above. Luminescence data were then analyzed to calculate the Z’-factor, to evaluate the reliability of the established assay for primary mAb screenings.

### R-ABA for high-throughput screening of mAb supernatants

The resazurin test plate was set-up as described above. After 2h incubation, 10 µL/well of resazurin solution were added and the samples further incubated for 2h before fluorescence measurement.

## RESULTS

### Development of the luminescent antibody bactericidal assay (L-ABA)

Accurate selection of the right source of complement to be used in bactericidal assays is crucial to correctly evaluate and interpret the functionality of antibacterial mAbs (14,15). Optimization of the amount of complement that can be used with a specific pathogen is equally important to ensure lack of toxicity and low non-specific killing of bacteria by complement alone. Baby rabbit complement (BRC) has been extensively used to evaluate the bactericidal potency of sera against *N. gonorrhoeae* and *N. meningitidis* (16–18). In addition, when compared to human complement, BRC is more easily available in large batches and subject to standardization (14). For all the above reasons, we decided to develop the L-ABA using BRC as the source of complement. The first steps consisted in optimizing the number of bacteria, the amount of BRC and incubation time to avoid complement toxicity and non-specific killing (NSK). Therefore, we incubated three different amounts of *N. gonorrhoeae* FA1090 (OD_600_ 0.1, 0.01 and 0.001) with four different percentages of BRC (5, 10, 20, 40%), for 1 or 2 hours at 37°C (**Fig. 2**).

**Fig. 2:**
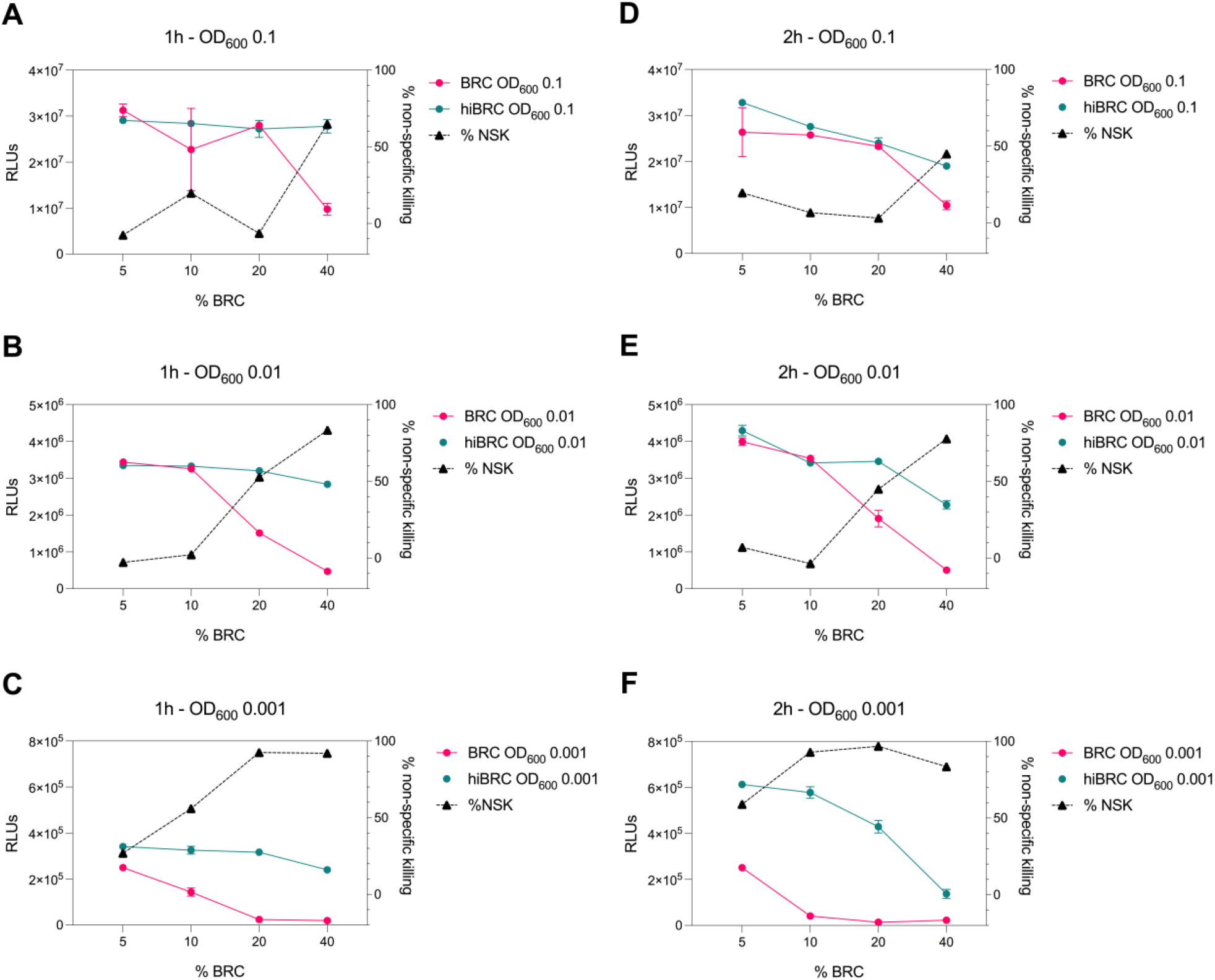
Evaluation of different conditions for L-ABA implementation. (A–C) Graphs show the viability and non-specific killing (NSK) of bacteria tested at OD_600_ 0.1 (A), 0.01 (B) and 0.001 (C) incubated with different amounts of BRC or hi-BRC (40, 20, 10, 5%) for 1h. (D – F) Graphs show the viability and non-specific killing of bacteria tested at OD_600_ 0.1 (D), 0.01 (E) and 0.001 (F) incubated with different amounts of BRC or hi-BRC (40, 20, 10, 5%) for 2h.

Bacterial viability was then evaluated by using the BacTiter Glo reagent as previously reported (19). This reagent measures the amount of ATP in the reaction which correlates with the number of viable microbial cells. Heat inactivated (hi) BRC was used as the control to evaluate the NSK induced by active BRC. Our data showed that 20% and 40% BRC caused the highest percentage of NSK in almost all tested conditions. Conversely, 5 and 10% of BRC, with the exception of 2h OD_600_ 0.001 condition, showed the least NSK. As for the number of bacteria, OD_600_ 0.1 and 0.01 resulted in the highest relative luminescence unit (RLU) values which are needed to correctly discriminate live from dead bacteria. Finally, both incubation times (1h and 2h) gave similar results, with the 2h condition resulting in higher RLUs at all ODs tested. Based on our data, four parameters were selected (10% BRC, 1h and 2h incubation, and OD_600_ 0.1 and 0.01) to evaluate the bactericidal efficacy of the known anti- *N. gonorrhoeae* monoclonal antibody 2C7 and determine the optimal settings for this assay (20). The antibody 2C7 showed complement-dependent killing of *N. gonorrhoeae* in all the tested conditions (**Fig. 3)**. However, OD_600_ 0.01 and 2h incubation resulted in the lowest half-maximal inhibitory concentration (IC_50_) value, suggesting higher assay sensitivity which is a desired feature to study the bactericidal activity of anti- *N. gonorrhoeae* mAbs in high throughput.

**Fig. 3:**
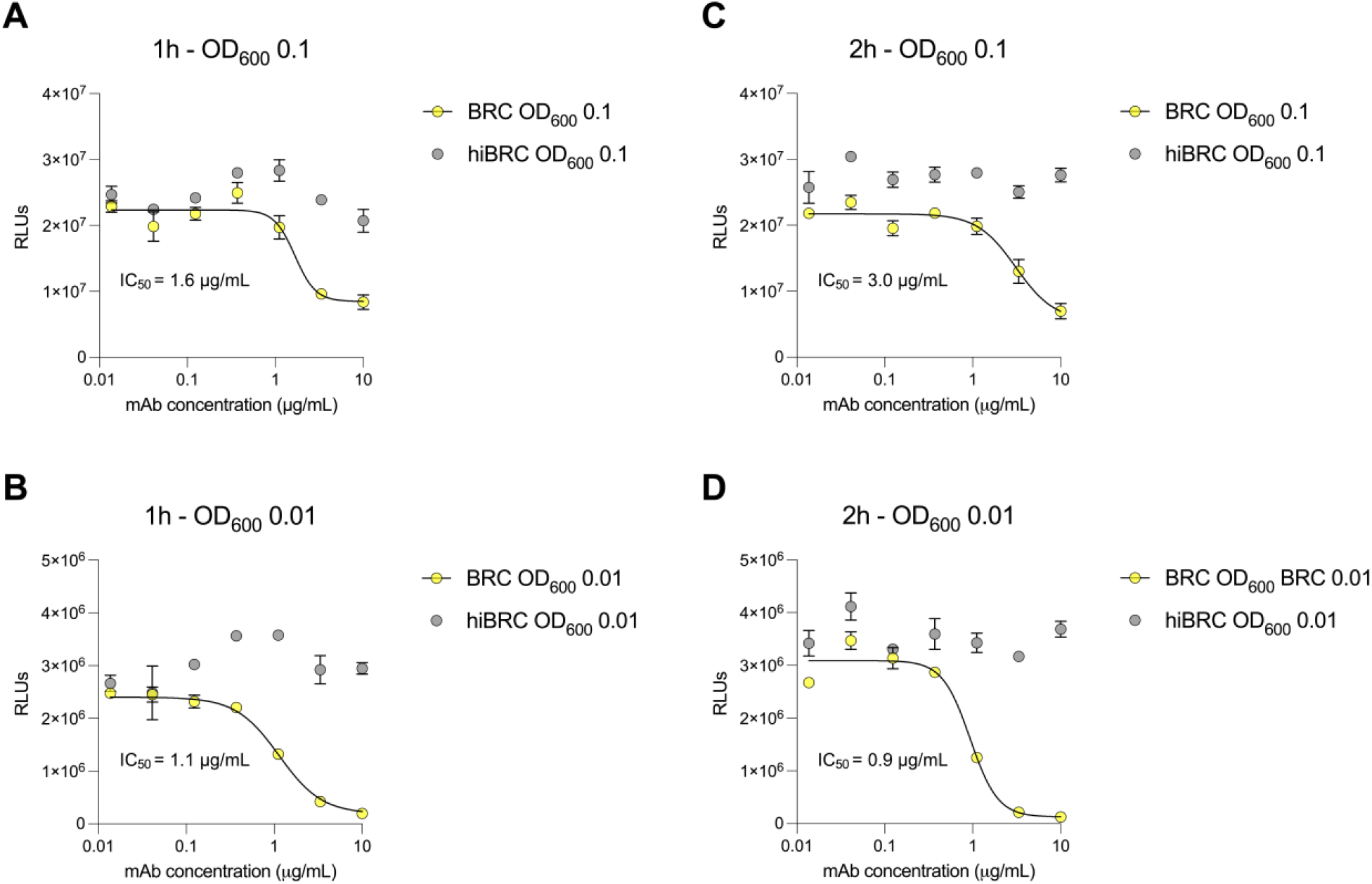
2C7 bactericidal activity evaluated by L-SBA. (A – B) Graphs show the bactericidal curves of 2C7 after 1h incubation with 10% BRC or hiBRC at OD_600_ 0.1 (A) and 0.01 (B). (C–D) Graphs show the bactericidal curves of 2C7 after 2h incubation with 10% BRC or hiBRC at OD_600_ 0.1 (C) and 0.01 (D). The IC_50_ values obtained with each condition are reported on the graphs.

### Correlation between L-ABA, R-ABA and C-SBA

The experimental conditions described above (OD_600_ 0.01, 10% BRC, 2h incubation) allowed to quantitatively evaluate the potency of the anti-gonococcal mAb 2C7 in terms of reduction of the ATP levels in the sample. To validate our assay, we compared the 2C7 bactericidal activity observed in L-ABA with a classical serum bactericidal assay (C-SBA), and a third approach that we named resazurin-based ABA (R-ABA). C-SBA relies on colony forming unit (CFU) counting and is conventionally used to evaluate serum activity. Differently, R-ABA is based on the reduction of the resazurin blue dye to a pink, fluorescent resorufin product by live bacteria. The amount of resorufin produced, and consequent fluorescent signal, in the assay is proportional to the number of live bacteria and to their metabolic activity. When we compared the IC_50_ obtained with the three assays, we observed almost identical values for L-ABA and R-ABA (0.25 and 0.23 µg/mL respectively), while a slightly lower IC_50_ was observed for the C-SBA (0.05 µg/mL) (**Fig. 4A-C**). The similarities among L-ABA, R-ABA and C-SBA are also demonstrated by a strong correlation among the three assays (r = 0.99, p < 0.001 L-ABA vs R-ABA and r = 0.83 p < 0.0039 L-ABA/R-ABA vs C-SBA) (**Fig. 4D-F**).

**Fig. 4:**
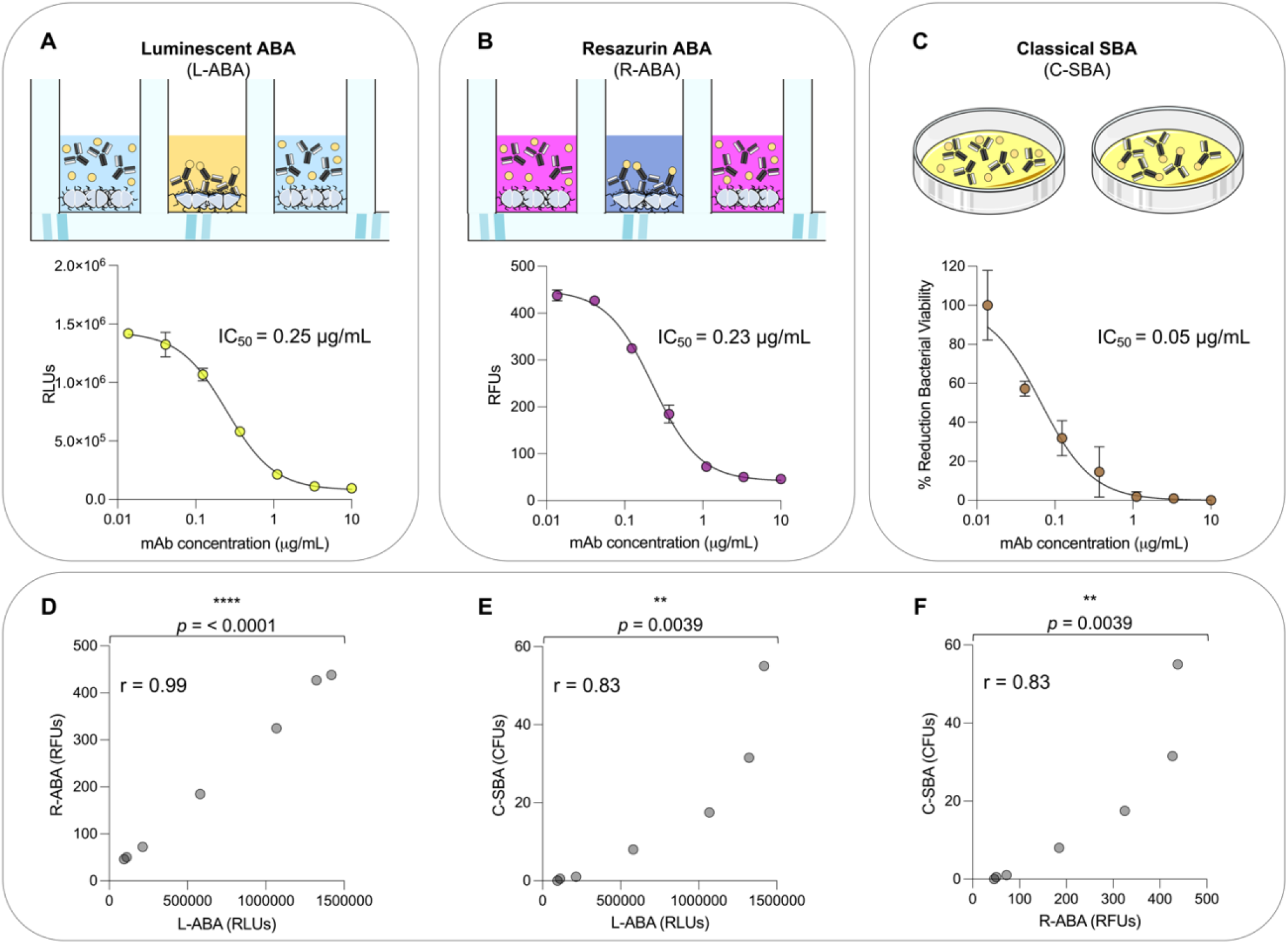
Comparison between L-ABA, R-ABA and C-SBA assays. (**A – C**) Bactericidal curves of 2C7 mAb starting from 10 µg/mL to 4 ng/mL (3-step serial dilutions) against *N. gonorrhoeae* FA1090 tested by L-SBA (A), R-SBA (B) and C-SBA (C). The IC_50_ values are reported on each graph. (D – F) Two-tailed Pearson coefficient correlation was performed between L-ABA and R-SBA (D), L-ABA and C-SBA (E) and R-ABA and C-SBA (F). The r squared values (r) are indicated on each graph. A nonparametric Mann–Whitney t test was used to evaluate statistical significances between groups. Two-tailed p-value significances are shown as **p < 0.01, and ****p < 0.0001.

### Implementation of high-throughput L-ABA (HT-L-ABA) and R-ABA (HT-R-ABA)

In high-throughput screenings, robust and sensitive assays are required to avoid false negative results and maximize the number of positive hits. This is particularly true in the case of primary screening, where hundreds or thousands of samples must be tested without the possibility to perform technical replicates. One of the most used parameters to assess the robustness and reliability of a primary screening assay is the Z factor, which is defined by two metrics: the mean (μ) and the standard deviation (σ) of both the positive (p) and negative (n) controls (21). Below, we report the formula used to calculate the Z factor:

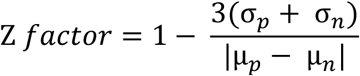

To mimic high-throughput mAb screening, small-scale 2C7 mAb expression in deep 96-well plates was obtained. This approach allowed to rapidly produce 48 independently expressed positive control replicates (2C7-containing Expi293F supernatants diluted 1:5) and 48 negative control replicates (mock transfections diluted 1:5) (**Fig. 5A**). The supernatants were used to implement high-throughput versions of our luminescence (HT-L-ABA) and fluorescence (HT-R-ABA) assays which aimed to assess in parallel all samples from a 96-well plate. The same experimental conditions previously described for the luminescence and fluorescence-based assays were applied to HT-L-ABA and HT-R-ABA. For both assays, all 48 wells containing 2C7 resulted in positive hits. Conversely, none of the mock transfection supernatants gave a false positive result. The experimental conditions adopted allowed us to reach a Z factor of 0.52 with HT-L-ABA and 0.68 with HT-R-ABA (**Fig. 5B-C**), indicating high robustness and specificity for both assays.

**Fig. 5:**
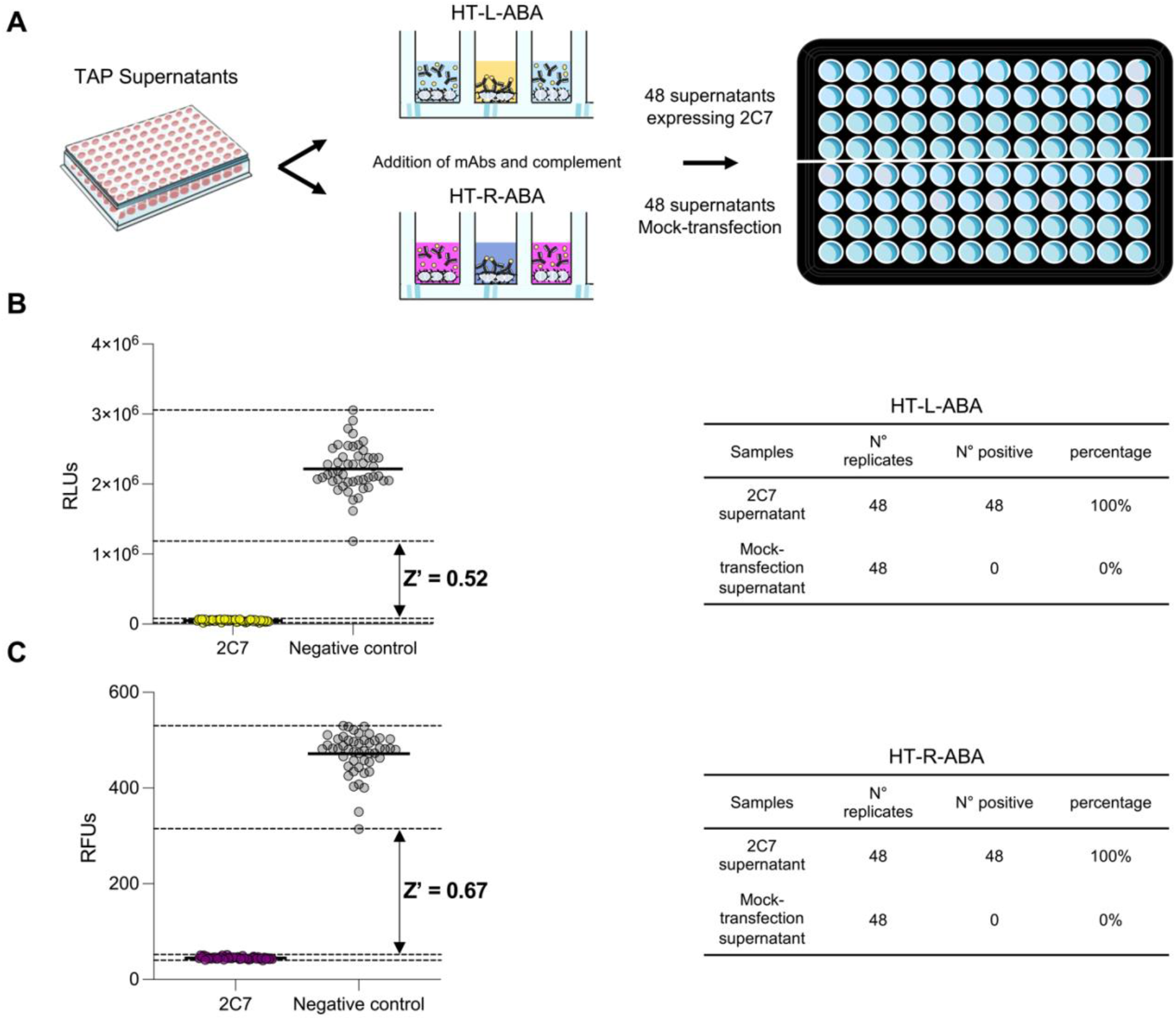
High-throughput assays for mAb screening. (A) Schematic representation of the two high-throughput assays (HT-L-ABA and HT-R-ABA) used to evaluate 2C7 bactericidal activity in a 96-well plate. (B) The graph shows the HT-L-ABA screening. The table describes number of replicates, positive hits, and percentage of positive hits for 2C7 supernatants and mock-transfected supernatants. (C) The graph shows the HT-R-ABA screening. The table describes number of replicates, positive hits, and percentage of positive hits for 2C7 supernatants and mock-transfected supernatants. Black lines represent the mean values, while black dotted lines display the maximum and minimum RLU values for both positive (2C7) and negative controls. The Z factor was calculated for both assays and is reported on the graphs.

## DISCUSSION

The global spread of multidrug-resistant gonorrhea has spurred efforts to develop safe and effective therapeutics and vaccines against this disease. Monoclonal antibodies have shown great potential as therapeutics and tools for antigen discovery and vaccine development against viral pathogens, while their use in the bacterial field is still limited. New assays are needed to implement high-throughput strategies to screen and identify highly bactericidal antibodies. In this paper we described the development of two new assays named L-ABA (high-throughput version HT-L-ABA) and R-ABA (high-throughput version HT-R-ABA). Differently from conventional assays, like the C-SBA, the methods presented in this work offer multiple advantages by eliminating the labor-intensive steps of plating hundreds or thousands of samples and counting individual CFU (2-4 h vs. >24 h), decreasing hands-on time by laboratory personnel and increasing the capacity to screen large sets of samples in a single day, thereby accelerating data acquisition and analyses. In addition, these approaches could also be applied to other *N. gonorrhoeae* strains, for rapid testing of new drugs, as well as other Gram-negative bacteria. Finally, the resazurin-based assay is an affordable method for mAb (or other drugs) screening, being resazurin not expensive, and results can be easily visualized by the naked eye. All these advantages can expedite the development of new therapeutic solutions and vaccines to fight *N. gonorrhoeae* and AMR.

## DATA AVAILABILITY STATEMENT

All data supporting the findings in this study are available within the article or can be obtained from the corresponding author upon request.

## AUTHOR CONTRIBUTIONS

S.S., M.T., E.A and R.R.: conceived the study. S.S. and M.T.: designed the experiments. S.S., M.T. and V.A.: conducted the experiments and analysed the data. S.S., M.T., E.A and R.R.: wrote the manuscript. S.S., M.T., V.A., C.S., E.A and R.R.: reviewed the manuscript.

